# Epithelium intrinsic zinc sensor controls immune homeostasis with gut microbes via regulation of Tuft cell lineage

**DOI:** 10.1101/2025.05.07.652635

**Authors:** Rebecca Yunker, Geongoo Han, Jorhenis Vasquez, Kellie Baldaro, Li Zhang, Sanghyun Lee, Shipra Vaishnava

## Abstract

Zinc is an essential micronutrient crucial for cell proliferation, differentiation, and apoptosis, yet its precise role in the constantly renewing intestinal epithelium remains unclear. We generated mice lacking the Zn-dependent transcription factor Metal-responsive Transcription Factor 1(MTF1^ΔIEC^) in intestinal epithelial cells. MTF1^ΔIEC^ mice exhibited altered metal homeostasis and acute susceptibility to Zn supplementation. Transcriptional and cellular analyses revealed increased inflammatory immune responses to microbes in MTF1^ΔIEC^ mice. Mechanistically MTF1 deletion resulted in the loss of Tuft cell lineages compromising barrier function against commensal microbes and pathogens. Ex vivo experiments demonstrated that at a cellular level, Zn treatment skewed the cellular composition from proliferating to differentiated cells. Specifically, we show that Zn sensing via MTF1 is required for IL-13 dependent induction of Tuft cells. Our findings underscore critical role of Zn in maintaining intestinal immune homeostasis through differentiation of specialized cell lineages, highlighting importance nutrient sensing in the constantly remodeling epithelial barrier.

## Introduction

The intestinal epithelium, a critical barrier between the host’s internal and external environments, plays a vital role in nutrient assimilation[1]. This dynamic tissue constantly renews and remodels itself in response to changing microbial and dietary cues, driven by stem cells at the base of intestinal crypts[2]. The intestinal stem cells produce progenitor cells that rapidly divide and differentiate into absorptive enterocytes and specialized secretory cells that are tasked with acquiring nutrients from the diet and defending against microbial encroachment[3, 4]. The lineage commitment of progenitors into the specialized cell types is intricately regulated by microbial, dietary, and immune factors, involving multiple signaling pathways, transcription factors, and epigenetic mechanisms [5]. While our understanding of this complex decision-making network is still evolving, the role of dietary factors has gained prominence, highlighting the evolutionary significance of micronutrients in driving intestinal epithelium regeneration[6–8].

Zinc (Zn) is a crucial dietary micronutrient that plays a vital role in maintaining intestinal homeostasis[9–11]. Approximately 10% of the human proteome relies on Zn, which is essential for enzyme function and transcription factor activity in processes like apoptosis, proliferation, and differentiation [12]. Zn deficiency and genetic mutations affecting Zn homeostasis have been linked to impaired intestinal barrier function and various gut-related disorders, including Acrodermatitis enteropathica and Crohn’s disease[13, 14]. While the importance of Zn in intestinal health is recognized, the specific molecular mechanisms by which Zn sensing in IECs impacts barrier formation and function, as well as host-microbiome interactions, remain poorly understood[15–17].

Cells have developed sophisticated mechanisms to regulate Zn uptake, distribution, and removal. IECs efficiently absorb dietary Zn and release it into circulation[18]. Metal response element-binding Transcription Factor-1 (MTF1) is a Zn-sensitive transcription factor which coordinates maintenance of intracellular Zn homeostasis by controlling the transcription of a various Zn-related machinery, including metallothioneins (MTs) and Zn-transporter-1 (ZnT1) [18]. MTF-1 senses Zn within the cell and coordinates the expression of these genes to help protect against oxidative stress, as well as metal toxicity [19]. MTs serve as labile pools of Zn which can readily accept or donate as necessary, acting as a Zn buffering system[20]. When MTF-1 is induced in a Zn-dependent manner, it regulates intracellular Zn levels by increasing the expression of MTs to chelate excess Zn and adjusting expression of Zn exporters [21, 22]. In this study we demonstrate that MTF-1 coordinates Zn homeostasis in IECs, and how its deletion leads to compromised barrier function, microbial encroachment, and inflammation, highlighting the intricate relationship between Zn homeostasis and intestinal health at a cellular level.

## Results

### MTF-1 regulates Zn homeostasis in the intestinal epithelium

Metal response element-binding Transcription Factor-1 (MTF1) serves as a critical Zn-sensing transcription factor in mammalian cells, orchestrating a complex network of Zn transport and storage proteins (Figure S1A)[21]. To investigate Zn sensing mechanisms in the intestinal epithelium, we generated mice with intestinal epithelial cell (IEC)-specific deletion of MTF1 (MTF1^ΔIEC^) using the Villin-Cre system (Fig. 1A). The efficiency of MTF1 deletion was confirmed by analyzing MT-2 expression, a specific MTF1 target in gut epithelium[23]. MTF1^ΔIEC^ mice showed significantly reduced MT-2 gene expression compared to MTF1^fl/fl^ controls as measured by qPCR (Fig. 1B). Immunofluorescence microscopy further validated this finding, revealing a marked reduction in MT protein expression within the intestinal crypts of MTF1^ΔIEC^ mice compared to the MT-rich crypts in control mice (Fig. 1C-D). MTF1 deletion affected the expression of various Zn transporters. Expressions of several Zn-related genes, including *zip3, znt1, znt2, and znt4,* were significantly downregulated in the intestine from MTF1^ΔIEC^ mice (Supplementary Fig. 1B-F), indicating comprehensive dysregulation of Zn transport machinery.

**Figure 1.**
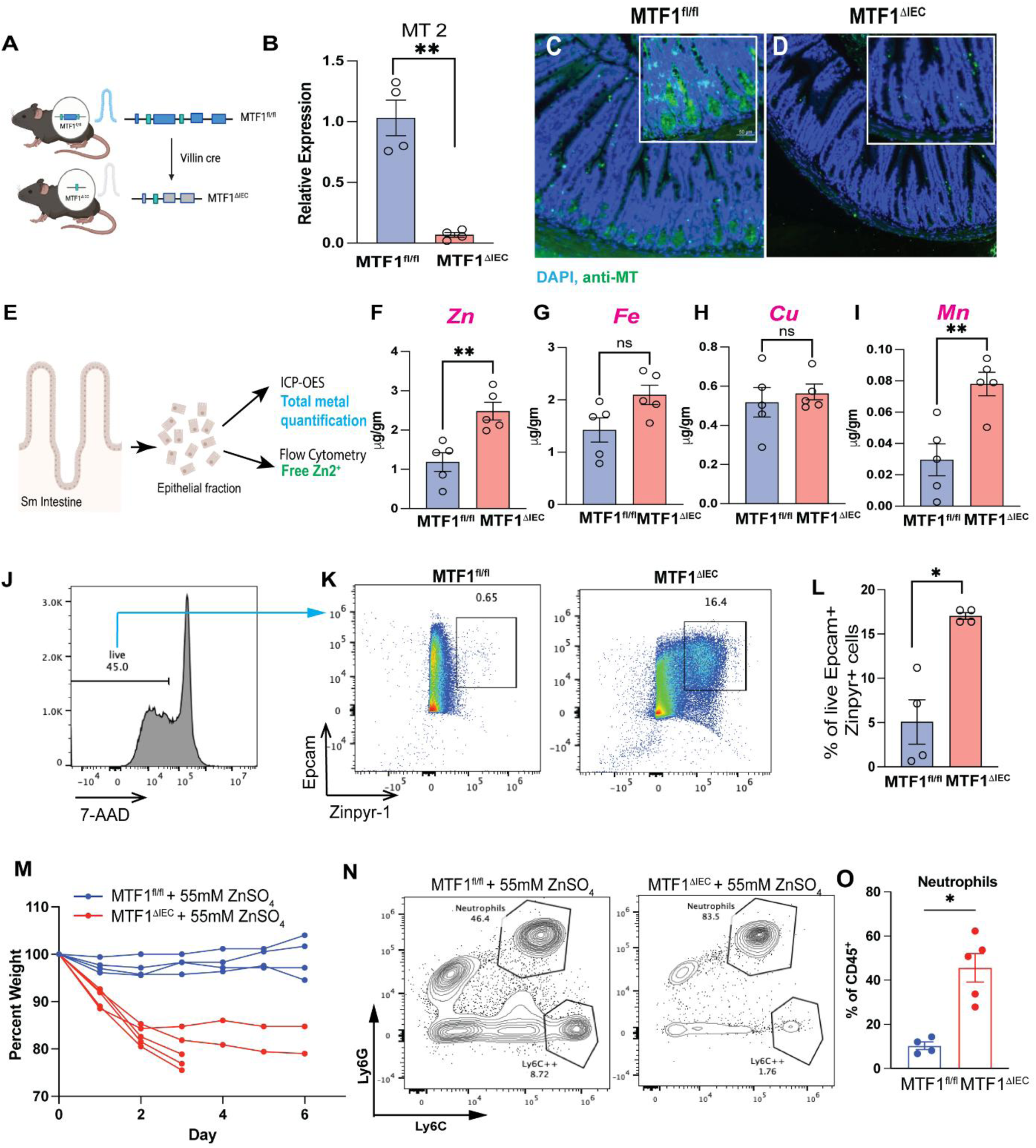
MTF-1 directs IEC intracellular metal homeostasis. A. Mouse model with intestinal epithelium-specific deletion of MTF1, generated using Cre-Lox recombination. B. Gene expression of metallothionein-2 (MT2) by qPCR (n=4 for each group). C. Immunofluorescent staining of MT1 and MT2 (green) proteins with nuclear DAPI stain (blue). E. Schematic demonstrating isolation of SI epithelial layer which was used in either ICP-OES for total metal quantification or flow cytometry for observing free Zn. F-I. ICP-OES quantification of IEC total (bound and free) displaying levels of manganese, Zn, copper and iron in MTF1^ΔIEC^ compared to littermate controls(n=5). J-L. Flow cytometry analysis of isolated IECs, which were stained with zinpyr-1, a cell-permeable fluorescent chelator of free Zn in cells(n=4). M. Mice were treated with 55mM ZnSO_4_ in drinking water for 1 week, with weights measured daily. Mice were euthanized when 20% bodyweight was lost (n=5). N-O. Mice were treated with 55mM ZnSO_4_ in drinking water for 4 days, and circulating lymphocytes were observed in blood by flow cytometry (n=5).

To assess the impact of MTF1 deletion on metal homeostasis, we performed comprehensive metal analysis using Inductively Coupled Plasma-Optical Emission Spectroscopy (ICP-OES) on the IECs isolated from the small intestine of MTF1^ΔIEC^ and MTF1^fl/fl^ littermates. This analysis revealed significant alterations in metal content within the epithelial fraction (Fig. 1E). Specifically, total Zn levels were significantly elevated in MTF1^ΔIEC^ IECs compared to controls (Fig. 1F). While iron and copper levels remained unchanged (Fig. 1G-H), manganese (Mn) showed a significant increase in MTF1^ΔIEC^ IECs (Fig. 1I). Mn and Zn transport in mammalian cells share significant overlap, primarily through transporters ZIP8, ZIP14, and ZnT10. These proteins, initially characterized for Zn transport, also play crucial roles in Mn homeostasis. This shared machinery leads to competitive transport between the metals and common regulatory mechanisms. To confirm changes were IEC specific and did not impact other organs, we analyzed metal contents of the liver by ICP-OES. The metal content in the liver did not show any significant changes indicating that the effect of MTF1 deletion in IECs on metal homeostasis is restricted to the intestinal epithelium (Fig. S1G). To specifically examine free Zn levels, we employed the Zn-specific fluorescent probe Zinpyr-1 in conjunction with flow cytometry. After gating on viable IECs (Fig. 1J), we observed a substantially higher percentage of Zinpyr-1-positive epithelial cells in MTF1^ΔIEC^ mice compared to controls, with quantification showing a significant increase in Zinpyr-1-positive cells (Fig. 1K and L). Additionally, MTF1^ΔIEC^ mice were intolerant to Zn supplementation in drinking water and showed significant weight loss compared to MTF1^fl/fl^ littermates when given 55mM ZnSO_4_ in drinking water (Fig. 1M). In response to Zn excess, MTF1^ΔIEC^ mice have significant increase in neutrophils in the peripheral blood at day 4 of treatment (Fig. 1N and O) indicating severe loss of barrier function. Together, these results reveal that the deletion of MTF-1 in IECs brings about distinct changes in expression of Zn machinery, leading to altered metal homeostasis in the IECs and reduced tolerance to Zn excess.

### Mice deleted for MTF1 in IECs have heightened inflammatory immune response and leaky barrier

To understand the impact of MTF1 deletion and disrupted Zn homeostasis in the epithelial cells on the intestinal tissue, we performed bulk RNA-sequencing on the ileum of MTF1^ΔIEC^ and MTF1^fl/fl^ littermates. Principal component analysis (PCA) of the data revealed that transcriptional response in ileum of MTF1^ΔIEC^ mice was significantly different than in MTF1^fl/fl^ littermates (Fig. S2A) with 24411 gene differentially expressed between the two groups of mice (Fig. S2B). Gene Ontology (GO) enrichment analysis identified key biological processes represented by the differentially expressed genes. The results GO enrichment analysis revealed that the ileum of MTF1^ΔIEC^ mice were significantly enriched in biological processes that were associated with upregulation of inflammatory immune response to microbes (Fig. S2C). In contrast the ileum of MTF1^ΔIEC^ mice showed a significant reduction in biological processes such as extracellular matrix organization, angiogenesis, and metal ion transport (Fig. S2D). Specifically, our data revealed that genes involved in inflammatory immune response to microbes are the most significantly upregulated change in small intestine of MTF1^ΔIEC^ compared to MTF1^fl/fl^ littermates (Fig. 2A). Expression of several inflammation-related genes such as *lcn2, il17a, lbp, tnf*, and *s100a* were significantly upregulated in small intestine along with genes involved in innate immune sensing of bacteria (*tlr2, nod1,* and *nod2*) in MTF1^ΔIEC^ compared to MTF1^fl/fl^ littermates (Fig. 2A). In accordance with RNAseq data, immune cell profiling of small intestinal tissue using flow cytometry revealed that MTF1^ΔIEC^ mice had significant alterations in inflammatory cell composition. MTF1^ΔIEC^ mice showed markedly increased neutrophil infiltration and tissue-resident macrophages (F4/80^+^CD64^+^) compared to control mice (Fig. 2B,C). Furthermore, examination of T cell populations revealed an enhanced Th17 cells were significantly enhanced in MTF1^ΔIEC^ mice. (Fig. 2D). These elevated immune responses suggest an augmented pro-inflammatory environment in the intestinal tissue of MTF1^ΔIEC^ mice.

**Figure 2.**
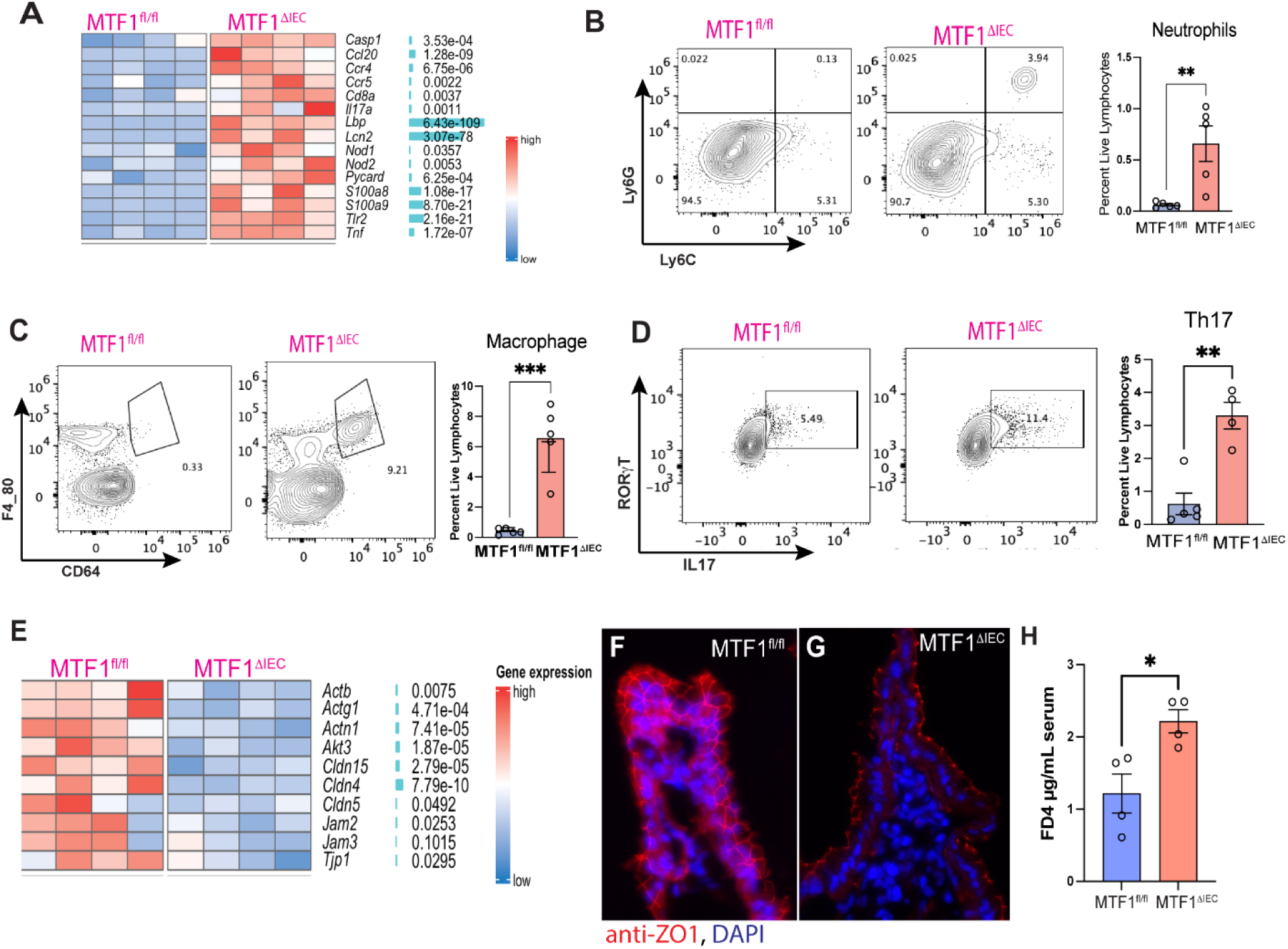
MTF1^ΔIEC^ mice show increased inflammatory immune response and a leaky intestinal barrier. A. RNA sequencing analysis of the terminal ileum reveals major changes in genomic expression of inflammatory markers in MTF1^ΔIEC^ compared to MTF1^fl/fl^ (n=4). B. Representative flow cytometry plot and quantification of neutrophils in epithelial fraction of terminal half of small intestine (SI) (n=5). C. Flow cytometry analysis of macrophages isolated from epithelial fraction of terminal half of SI (n=5). D. Flow cytometry analysis of Th17 cells in the lamina propria of terminal half of SI (n=4). E. RNA sequencing analysis of terminal ileum shows decrease in tight–junction related proteins in MTF1^ΔIEC^ compared to MTF1^fl/fl^ (n=4). F-G. Immunofluorescent staining of Zonula Occludens-1 (ZO1) in SI of MTF1^ΔIEC^ compared to MTF1^fl/fl^. H. Mice were given 6mg/gram bodyweight FITC-dextran 4 kDa (FD4) by oral gavage, 3 hours before mouse was euthanized, to test intestinal permeability. Graph shows concentration of FD4 in serum of each group.

Concurrent with an increase in inflammatory immune cells, MTF1^ΔIEC^ mice showed a reduction in tight junctions and adhesion molecules which maintain intestinal barrier integrity (Fig. 2E). The small intestine from MTF1^ΔIEC^ showed a significant decrease in Zonula Occludens-1 (ZO1), which localizes at tight junctions between cells (Fig. 2F,G). Barrier functionality was assessed by administering 4 kDa FITC-dextran (FD4) via oral gavage, followed by measurement of the fluorescence signal in the serum. As a result, MTF1^ΔIEC^ mice showed an increased barrier permeability with higher FD4 levels compared to control (Fig. 2H). These findings collectively demonstrate that loss of Zn sensing in IECs results in loss of immune homeostasis characterized by upregulated inflammatory gene expression, increased recruitment of specific innate and adaptive immune cells, and a leaky barrier in the intestinal tissue.

### MTF1^ΔIEC^ mice show a diminished numbers of Paneth Cells and loss of spatial segregation with gut microbiome

The intestinal epithelium comprises specialized cells that play a critical role in maintaining immune homeostasis by regulating interactions with gut bacteria[24]. To assess if loss of Zn homeostasis in IECs due to MTF1 deletion affects the specialized secretory cells that are key for regulating barrier function against gut microbes, we used bulk RNA-sequencing data from small intestinal tissues of MTF1^ΔIEC^ and MTF1^fl/fl^. We used a cell marker gene list for specific epithelial cell types curated from publicly available scRNA data sets such as PanglaoDB[25]. Our analysis revealed that expression of Tuft cell (Padj=2.53E^09^, Fig.3A) and Paneth cell (Padj=0.0003, Fig.3A) specific gene sets were significantly downregulated in small intestinal tissue of MTF1^ΔIEC^ compared to MTF1^fl/fl^ littermate controls. No significant changes were observed in Goblet cell markers (Fig.3A). Gene expression of another rare secretory epithelial cell lineage called enteroendocrine cells that secretes gut hormones and aids in food digestion and absorption did not change significantly (Fig.3A).

**Figure 3.**
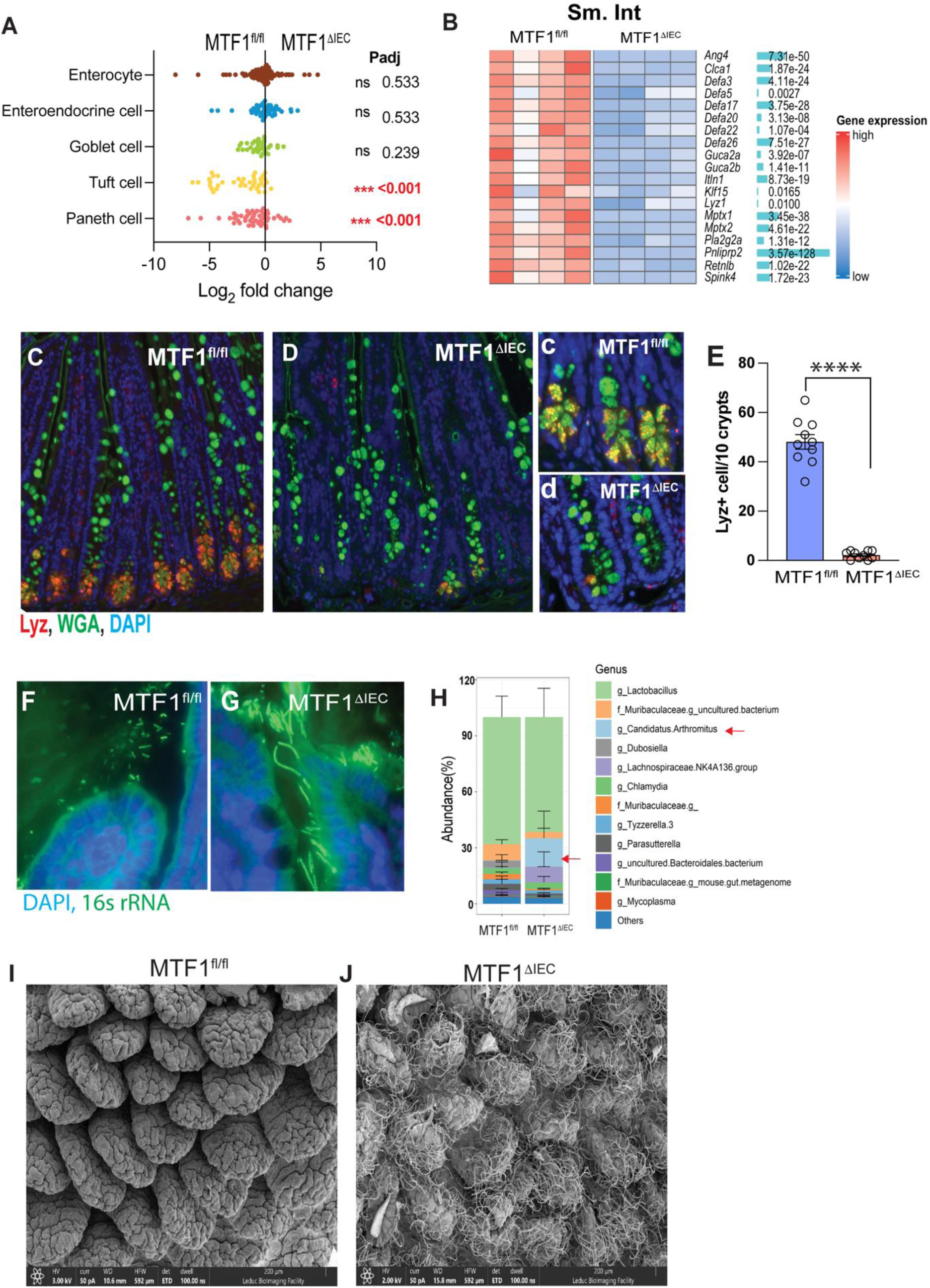
MTF^ΔIEC^ mice display fewer Paneth Cells concurrent with increased mucosa associated bacteria. A. Bulk RNA-sequencing results of specific epithelial cell types in intestine of MTF1^ΔIEC^ and MTF1^fl/fl^ littermate controls. B. RNA-sequencing results of Paneth cell-markers (n=4). C-D. MTF1^ΔIEC^ and MTF1^fl/fl^ SI sections stained with lysozyme (red) lectins (green) and DAPI (blue). E. Numbers of Paneth cells per 10 crypts counted. F-G. Fluorescent In Situ Hybridization (FISH) staining of mucosal adherent bacteria. H. Comparison of relative abundance of bacteria at family and genus level of mucus scraping (n=4). J. Scanning Electron Microscopy (SEM) images of terminal ileum villi.

RNA-sequencing results showed a broad decrease in Paneth cell-related markers, such as the family of alpha-defensin antimicrobials and MMP7, an enzyme present in Paneth cells involved in the activation of alpha-defensins (Fig. 3B). Immunofluorescence assay with anti-Lysozyme (Lyz) antibody(red) that mark the Paneth cells in addition to wheat germ agglutinin (WGA) and ulex europaeus agglutinin (UEA-I) which stains mucus-producing cells such as goblet cells(green), revealed that Paneth cells are significantly reduced in the small intestine of MTF1^ΔIEC^ compared to MTF1^fl/fl^ (Fig. 3C-E). In contrast, mucus secreting goblet cells were present in both consistent with the RNA-sequencing data showing significant reduction in Paneth cell markers but not Goblet cells in small intestinal tissue of MTF1^ΔIEC^ compared to MTF1^fl/fl^ (Fig. 3A).

Paneth cell antimicrobial response is key for regulating interactions with gut microbes. The antimicrobial peptides secreted by Paneth cells, particularly α-defensins, limit the number of commensal bacteria attaching to the mucosa and help maintain immune homeostasis. Microbial dysbiosis characterized by increased adherent microbes such as segmented filamentous bacteria (SFB) is observed in case of defective antimicrobial production by Paneth cells [26]. We assessed if the loss of Paneth cells in MTF1^ΔIEC^ mice coincided with microbial dysbiosis, such as an increase in total bacterial load or defects in spatial segregation of gut bacteria from the host. We used fluorescent *in-situ* hybridization (FISH) analysis that specifically labels bacteria (16s rRNA probe). We observed that the epithelium adherent microbes are significantly enriched in the mucus of MTF1^ΔIEC^ mice compared to MTF1^fl/fl^ littermate controls (Fig. 3F-G). 16s qPCR analysis revealed that MTF1^ΔIEC^ had higher bacterial loads compared to MTF1^fl/fl^ littermate controls in SI (Fig. 3H). This change was reflected in 16S sequencing analysis of the bacterial communities in the mucosal contents in the ileum. SFB, also referred to as *Candidatus Arthromitus*, was significantly increased in the mucus of MTF1^ΔIEC^ mice, and other healthy bacteria were associated more strongly with control (Fig. 3H). Numerous SFB were observed tightly attached to the villi by Scanning Electron Microscopy (SEM), even after undergoing multiple flush and wash steps during preparation (Fig. 3I-J). Taken together our data show that MTF1 deletion in epithelial cells results in loss of functional Paneth cells that are key for regulating host-microbe interactions in the gut.

### MTF1^ΔIEC^ mice lack tuft cells, related immune circuitry, showing altered susceptibility to worm and viral infections

RNA-sequencing analysis revealed a significant reduction in tuft cell-specific genes in MTF1^ΔIEC^ mice compared to controls, including key markers *Il25, Dclk1,* and *Trpm5* (Fig. 4A). Immunofluorescence staining for the tuft cell marker DCLK1 confirmed a dramatic reduction in tuft cell numbers in MTF1^ΔIEC^ mice compared to MTF1^fl/fl^ controls (Fig. 4B-D). Reduction in tuft cells was seen throughout the length of small intestine (Fig. S3A and S3B). Comparable load of commensal protozoa *Tritrichomonas musculus,* which is known to induce tuft cells in laboratory mice, was found in MTF1^ΔIEC^ mice and MTF1^fl/fl^ controls (Fig. S3C), suggesting that loss of tuft cells in MTF1^ΔIEC^ mice is independent to tuft cell-stimulating microbes.

**Figure 4.**
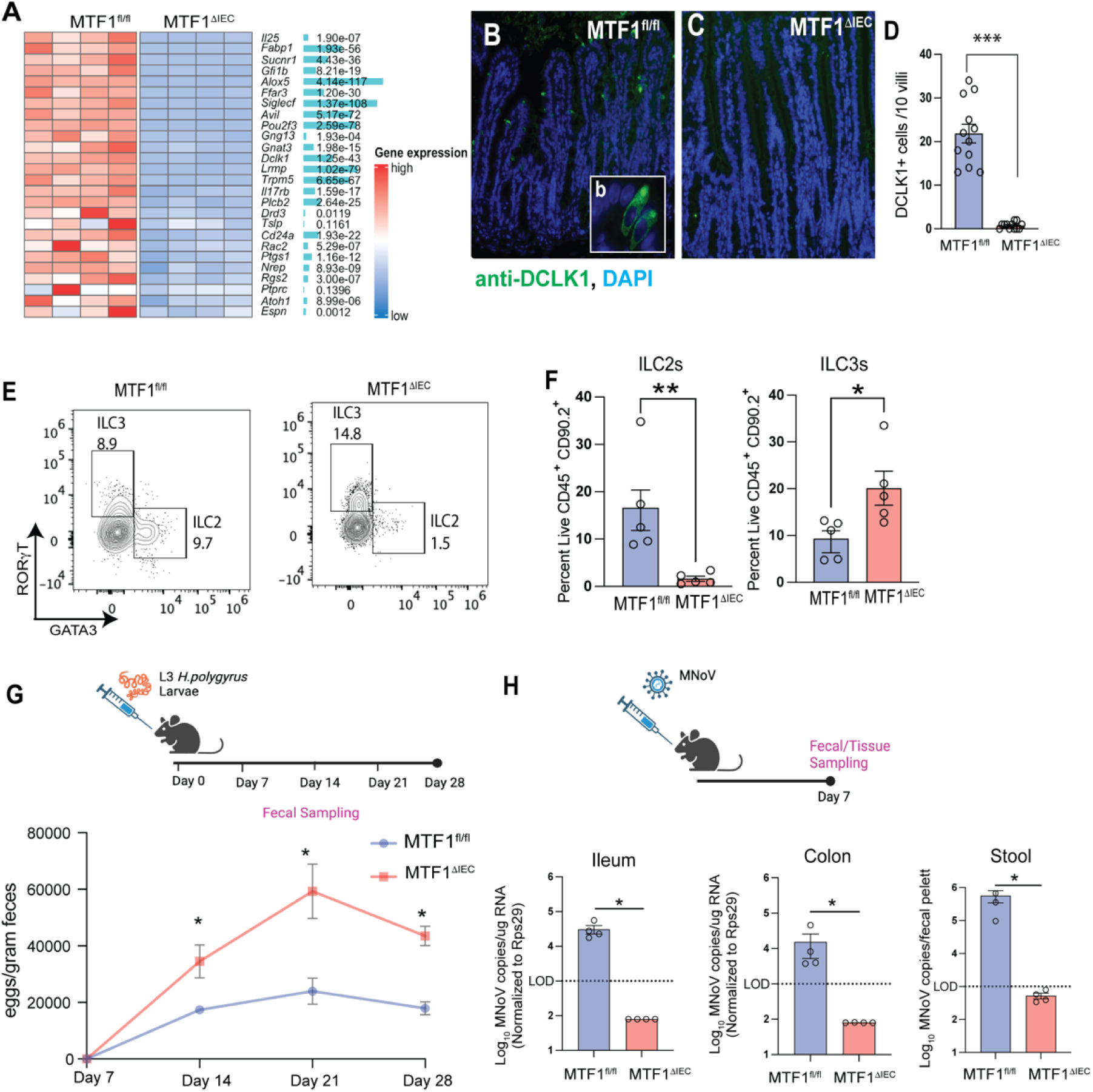
MTF1^ΔIEC^ mice lack tuft cells and tuft cell dependent immune circuitry leading to altered immunity. A. RNA-sequencing results of tuft cell-markers (n=4). B-C. Tuft cells (green) by immunofluorescent staining of DCLKM-1 (DCLK-1) and DAPI (blue). D. Number of tuft cells counted per 10 villi. E-F. Flow cytometry analysis of lymphocytes isolated from the lamina propria of terminal half of SI (n=5). G. Mice were treated with *H. polygyrus* larvae and egg burden was measured weekly (n=5). H. Mice were infected with murine norovirus (MNoV), and after 1-week intestinal tissues and fecal samples were harvested. MNoV copies were detected by qPCR (n=4). L.O.D., limit of detection.

Tuft cells are the exclusive source of IL-25 in the small intestine. IL-25 produced by tuft cells is a key activator of innate lymphoid cell (ILC)-2 in the intestine [27, 28]. Flow cytometry analysis revealed a significant alteration of ILC populations in MTF1^ΔIEC^ mice (Fig. 4E, S3D). The percentage of ILC2s was markedly reduced, while ILC3s showed a significant increase in the SI of MTF1^ΔIEC^ mice compared to MTF1^fl/fl^ controls (Fig. 4E-F).

Tuft cells play a crucial role in the immune response against *Heligmosomoides polygyrus* infection in the small intestine [27]. To assess functional consequences of tuft cell loss, we challenged MTF1^ΔIEC^ and MTF1^fl/fl^ mice with *H. polygyrus*. MTF1^ΔIEC^ mice showed impaired helminth control, with significantly higher fecal egg burdens emerging by day 14 post-infection and persisting through day 28 compared to MTF1^fl/fl^ controls (Fig. 4G).

Tuft cells are essential for murine norovirus (MNoV) to establish persistent infection in the intestine [29]. Tuft cell-deficient mice cannot support chronic viral shedding in feces, ileum, and colon. We next examined susceptibility to murine norovirus (MNoV). One week after infection, MTF1^ΔIEC^ mice showed significantly reduced viral loads in ileum, colon, and stool samples compared to controls, with levels falling below the limit of detection (Fig. 4H). Taken together our data show that MTF1 deletion in epithelial cells results in loss of tuft cells at a transcriptional, cellular and functional level.

### Induction of MTF1 deletion in adult mice leads to rapid tuft cell loss and followed by subsequent defects in Paneth cells

Paneth cell biology is uniquely dependent on Zn, where Zn chelation or loss of Zn transporter results in Paneth cell atrophy[30–35]. Recent studies have further suggested there is bidirectional communication between Paneth and tuft cells[36, 37]. To determine if disruption of Zn sensing in MTF1^ΔIEC^ mice results in concurrent or sequential loss of Paneth cell and Tuft cell lineages, we generated an inducible IEC-specific MTF1 deletion mouse model (MTF1^ERTΔIEC^) and littermate control by crossing MTF1^fl/fl^ mice with Vill-creERT2 strain. Adult mice at 8 weeks of age were treated with tamoxifen for 5 days to induce MTF1 deletion. Mice were observed at either 2 or 4 weeks after the induction of MTF1 deletion(Fig. 5A, F). At 2 weeks, MTF1^ERTΔIEC^ mice showed significant loss of tuft cell markers by qPCR (Fig. 5B). Immunofluorescent staining further revealed a dramatic loss of DCLK1^+^ tuft cells in MTF1^ERTΔIEC^ mice compared to littermate controls (Fig. 5C). However, Paneth cell numbers remained largely unchanged suggesting that loss of Zn sensing results in rapid loss of tuft cells but not Paneth cell lineages at the early time point (Fig. 5D-E). At 4 weeks post-induction, we observed a partial loss of Paneth cells markers were observed such as antimicrobial factors like lysozyme, MMP7, and defensin10 (Fig. 5I-J). Immunofluorescent staining shows reduction in lysozyme signal in the crypts of MTF1^ERTΔIEC^ mice (Fig. J). Paneth cells have been shown to remain longer in the epithelium, persisting up to 6-8 weeks, therefor it is possible that Paneth cell defect appear later compared to tuft cell [38] [39]. We continued to observe a significant reduction in tuft cell markers by qPCR and immunofluorescent staining at the 4 weeks post-induction time frame suggesting that loss Zn sensing in IECs first results in rapid loss of Tuft cell lineage and a subsequent defect in Paneth cell lineage at a later time point (Fig. 5G-H).

**Figure 5.**
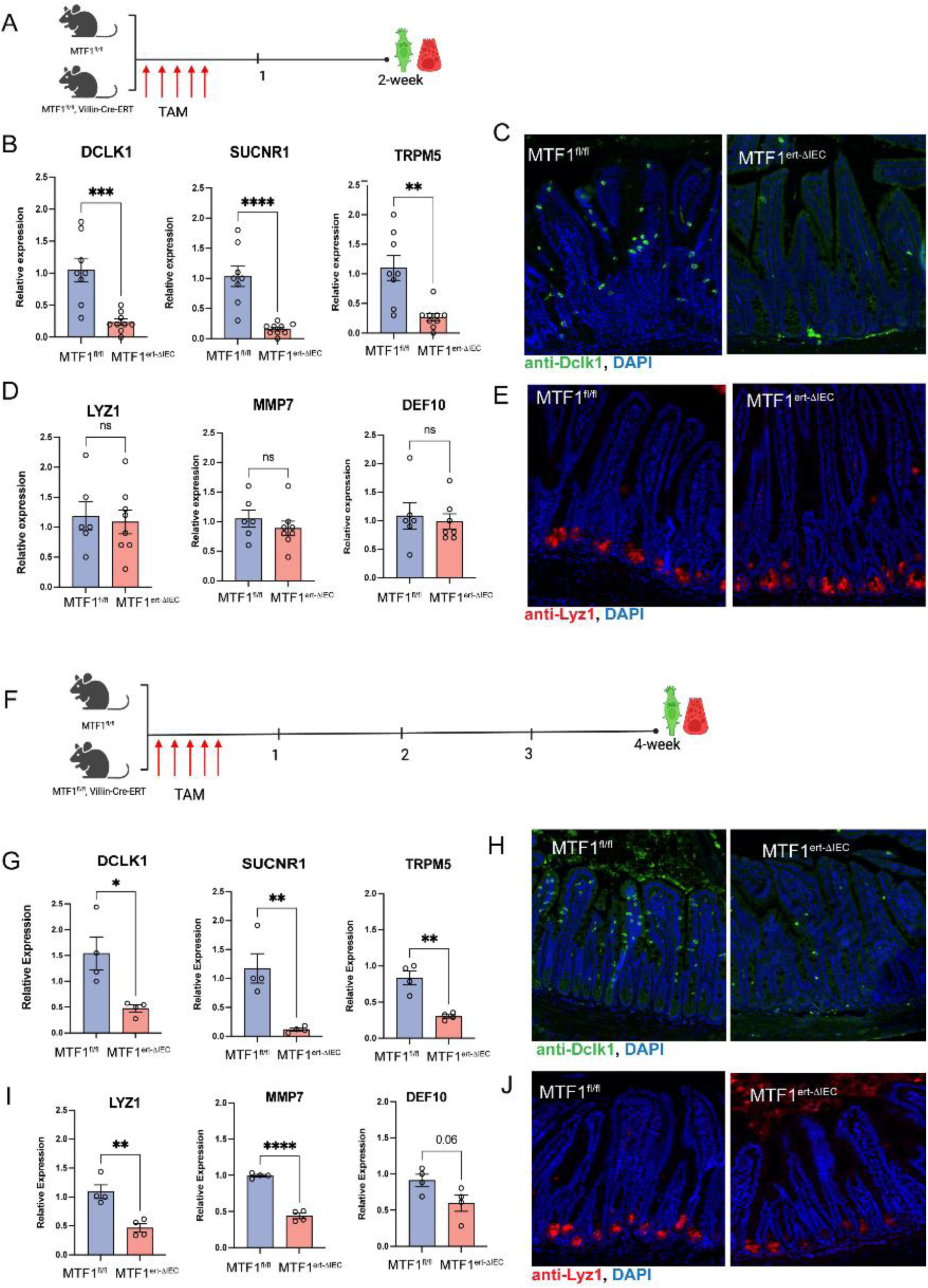
Tuft cells are rapidly lost after the induction of MTF1 deletion whereas Paneth cell defect appears later. A,F. Mice were given tamoxifen at 50mg/kg by intraperitoneal injection for 5 days to induce deletion on MTF1 by Cre-ERT2. Mice were observed at 2- and 4-weeks post induction. B. Tuft cell markers were observed by qPCR after 2 weeks of Cre induction in terminal SI (n=8-9). C. Tuft cells (green) by immunofluorescent staining of DCLKM-1 (DCLK-1) and DAPI (blue) in MTF1^ERTΔIEC^ mice compared to MTF1^fl/fl^ terminal SI. D. Paneth cell markers observed by qPCR in terminal SI. E. Immunofluorescent staining terminal SI of Paneth cells stained with lysozyme (red) and DAPI (blue). G. Tuft cell markers were observed by qPCR 4 weeks post-induction (n=4). H. Tuft cells (green) by immunofluorescent staining of DCLKM-1 (DCLK-1) and DAPI (blue) in MTF1^ERTΔIEC^ mice compared to MTF1^fl/fl^ terminal SI. I. Paneth cell markers observed by qPCR in terminal SI. J. Immunofluorescent staining terminal SI of Paneth cells stained with lysozyme (red) and DAPI (blue).

### MTF1 mediates Zn-dependent switch from proliferation to differentiation in intestinal epithelial cells ex vivo

Zn is recognized as a crucial factor for growth and proliferation in mammalian cells, yet how this dietary micronutrient affects intestinal epithelial cell proliferation and differentiation is not known. Studies have shown that continuously dividing cells are highly sensitive to intracellular level of Zn [40, 41]. Optimal level of Zn^2+^ ions are needed for dividing cells to exit cell cycle, and excess or insufficient intracellular Zn^2+^ can stall cells in the cell cycle. Given that MTF1 deletion results in mismanagement of Zn in IECs, we asked whether intestinal organoids derived from MTF1^ΔIEC^ and MTF1^fl/fl^ show differences in proliferation and differentiation of IECs [42]. Bulk RNA-sequencing revealed small intestinal organoids (SIOs) from MTF1^ΔIEC^ had significantly less expression of tuft cell and enterocytes specific genes during homeostasis, however no significant difference was observed in expression of Paneth cell-specific genes after 7-days of growth (Fig. 6A, Fig. S5E, and Fig. S5F). Due to the absence of Zn in standard serum-free organoid media (Fig. S4A), supplementation of Zn was used to bring cellular zinc levels to physiological levels and showed minimal impact on organoid budding (Fig. S4B). Bulk RNA-sequencing of the organoids revealed that Zn treatment resulted in enhancing the difference between MTF1^ΔIEC^ and MTF1^fl/fl^ SIOs (Fig. 6A-B). Upon Zn treatment, MTF1^ΔIEC^ SIOs have a significant decrease in markers for differentiated absorptive and secretory epithelial cell lineages such as enterocytes, enteroendocrine and tuft cells compared to Zn-treated MTF1^fl/fl^ SIOs (Fig. 6B). GO enrichment analysis further corroborated that MTF1^ΔIEC^ SIOs showed a decreased ability to perform fatty acid metabolism, small molecule metabolism and intestinal absorption, actions often performed by differentiated cell types such as enterocytes (Fig 6C). Instead, MTF1^ΔIEC^ SIOs showed an increase in genes associated with cell proliferation such DNA replication, nuclear division, and chromosome segregation (Fig 6D).

**Figure 6.**
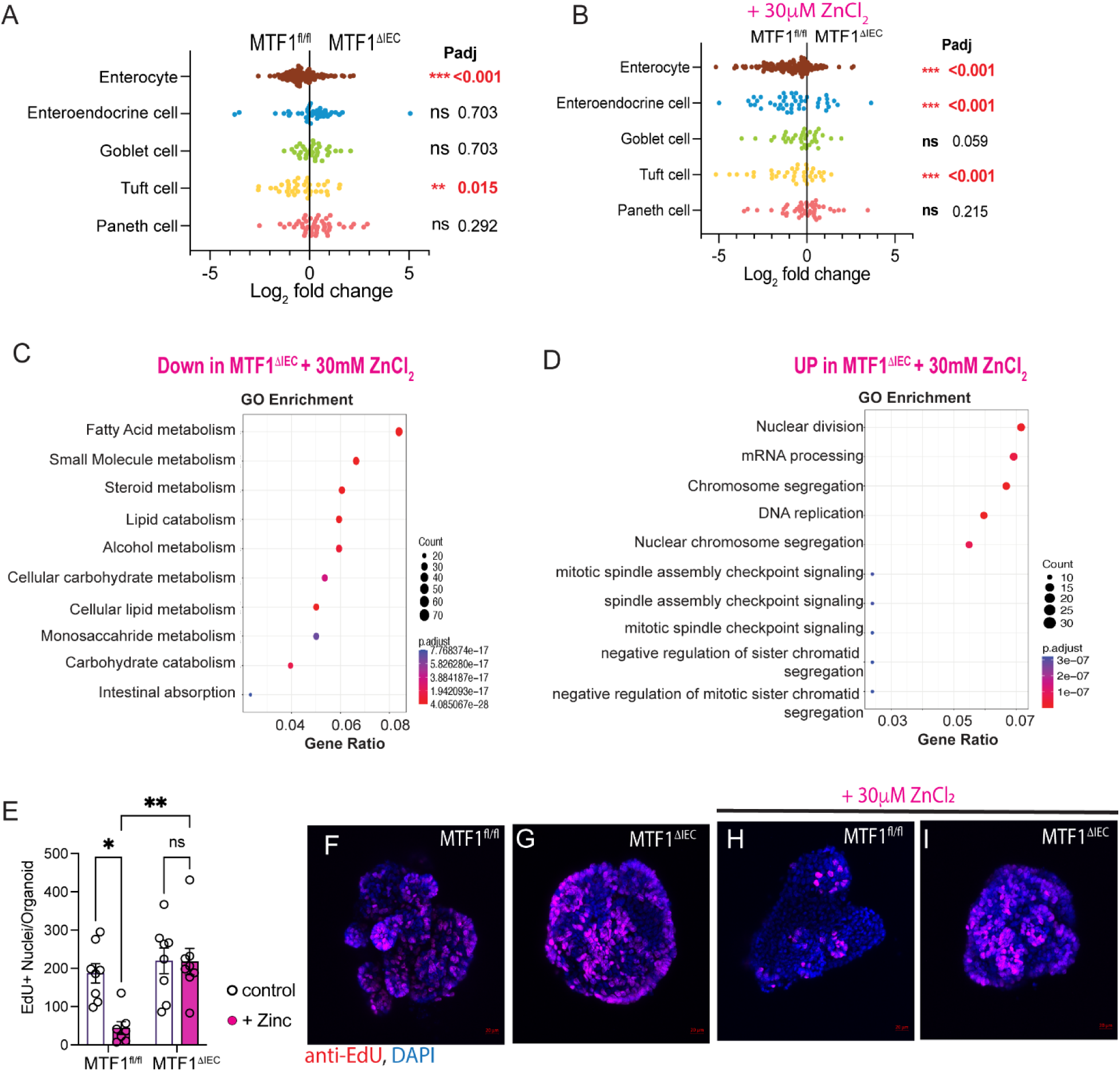
MTF1 is required for Zn-dependent reduction in proliferation ex vivo. Organoids were grown with for 7 days in homeostasis or with 30µM ZnCl_2_ present in media. A. **Bulk** RNA-sequencing results show changes in specialized cells in MTF1^ΔIEC^ in homeostasis. B. Comparison of MTF1^ΔIEC^ and MTF1^fl/fl^ SIOs with 30µM ZnCl_2_ treatment, showing genes associated with differentiated IECs. C. GO enrichment analysis of genes downregulated in MTF1^ΔIEC^ SIOs with 30µM Zn treatment. D. GO enrichment analysis of genes upregulated in MTF1^ΔIEC^ SIOs after 30µM Zn treatment. E. EdU incorporation into SIOs occurred for 2 hours before harvest. E. EdU positive nuclei were counted per organoid, as observed from whole-mount staining using EdU (pink) and DAPI (blue). F-G. Representative images of EdU staining in MTF1^fl/fl^ and MTF1^ΔIEC^ SIOs without Zn treatment. H-I. Representative images of EdU staining in MTF1^ΔIEC^ and MTF1^fl/fl^ SIOs with Zn treatment.

We did not observe as significant of a reduction in Paneth cell marker gene set between MTF1^ΔIEC^ and MTF1^fl/fl^ SIOs as we did *in vivo* (Fig. 6A). However fewer lysozyme positive cells were observed in MTF1^ΔIEC^ SIOs compared to MTF1^fl/fl^ SIOs (Fig. S5A). After the introduction of Zn, this difference between MTF1^ΔIEC^ and MTF1^fl/fl^ SIOs became even more significant, as Zn treated MTF1^fl/fl^ SIOs showed a significant increase in lysozyme-positive Paneth cells per organoid (Fig. S5B). Transcriptional analysis showed that Zn supplementation in MTF1^ΔIEC^ SIOs compared to MTF1^fl/fl^ SIOs, significantly reduced expression of some, but not all, of the genes essential for Paneth cell function, such as *Lyz1* and *Ang4* (Fig. S5C). The expression of genes involved in Paneth cell specification such as *Lgr4* (Leucine-rich repeat-containing G protein-coupled receptor 4) and *Ctnnb1* (β-catenin) are also significantly reduced in MTF1^ΔIEC^ SIOs upon Zn supplementation compared to MTF1^fl/fl^ SIOs (Fig. S5D).

To observe proliferating cells, MTF1^ΔIEC^ and MTF1^fl/fl^ SIOs with or without Zn treatment were labelled with EdU for 2 hours on day 7 of growth. Total number of EdU+ cells per organoid were similar between MTF1^ΔIEC^ and MTF1^fl/fl^ SIOs without Zn treatment (Fig. 6E). With Zn treatment, the number of Edu+ cells per organoid was significantly diminished in MTF1^fl/fl^ SIOs compared to MTF1^fl/fl^ SIOs without Zn treatment, whereas this effect was absent in MTF1^ΔIEC^ SIOs(Fig.6E). In Zn-treated MTF1^fl/fl^ SIOs, EdU+ cells were restricted to the crypt-like domain (Fig 6 H). In contrast, Zn treated-MTF1^ΔIEC^ SIOs exhibited EdU+ cells distributed all throughout the organoid, a pattern similar to that observed in untreated MTF1^ΔIEC^ and MTF1^fl/fl^ SIOs (Fig. 6F-I). Taken together our results demonstrate that MTF1 is crucial for the Zn-dependent switch from proliferation to differentiation in intestinal organoids.

### MTF1-dependent Zn homeostasis regulates tuft cell differentiation and maintenance upon IL-13 induction

Tuft cells are rare chemosensory cells in SIOs making tuft cell quantification challenging. To determine if loss of tuft cell lineages were due to cell intrinsic defects in Zn sensing, SIOs from MTF1^ΔIEC^ and MTF1^fl/fl^ were stimulated with IL-13 and DCLK1^+^ cells were quantified. Upon IL-13 stimulation, both MTF1^ΔIEC^ and MTF1^fl/fl^ SIOs showed increased numbers of tuft cells with no significant difference in tuft cells numbers (Fig. 7A-B, E) suggesting that stem cells were not defective in generating tuft cells. Next, we tested if addition of physiological level of Zn to the media would alter induction of tuft cells in MTF1^ΔIEC^ SIOs that show altered Zn sensing. Upon introduction of Zn in combination with IL-13 stimulus, MTF1^ΔIEC^ had significantly fewer tuft cells compared to MTF1^fl/fl^ SIOs (Fig. 7C-E). This demonstrates that in the presence of Zn, MTF1^ΔIEC^ SIOs that have a defect in Zn sensing, and cannot differentiate or retain tuft cells. Additionally, EdU was administered to SIOs 2 hours before harvest, to observe if ILl-13 stimulation with or without Zn treatment impacted proliferation. In IL-13 treatment, MTF1^ΔIEC^ SIOs showed reduced proliferating cells compared to controls (Fig. 7F). After combination of IL-13 and Zn treatment, MTF1^fl/fl^ SIOs exhibited a significant reduction in proliferation compared to MTF1^fl/fl^ SIOs treated with IL-13 alone. However in MTF1^ΔIEC^ SIOs, the addition of Zn had no effect on proliferation compared to IL-13 treatment alone. These results point to the importance of Zn sensing via MTF1 mediating differentiation of Tuft cells, where the alteration of Zn homeostasis upon MTF1 deletion results in loss of tuft cells *ex vivo*.

**Figure 7.**
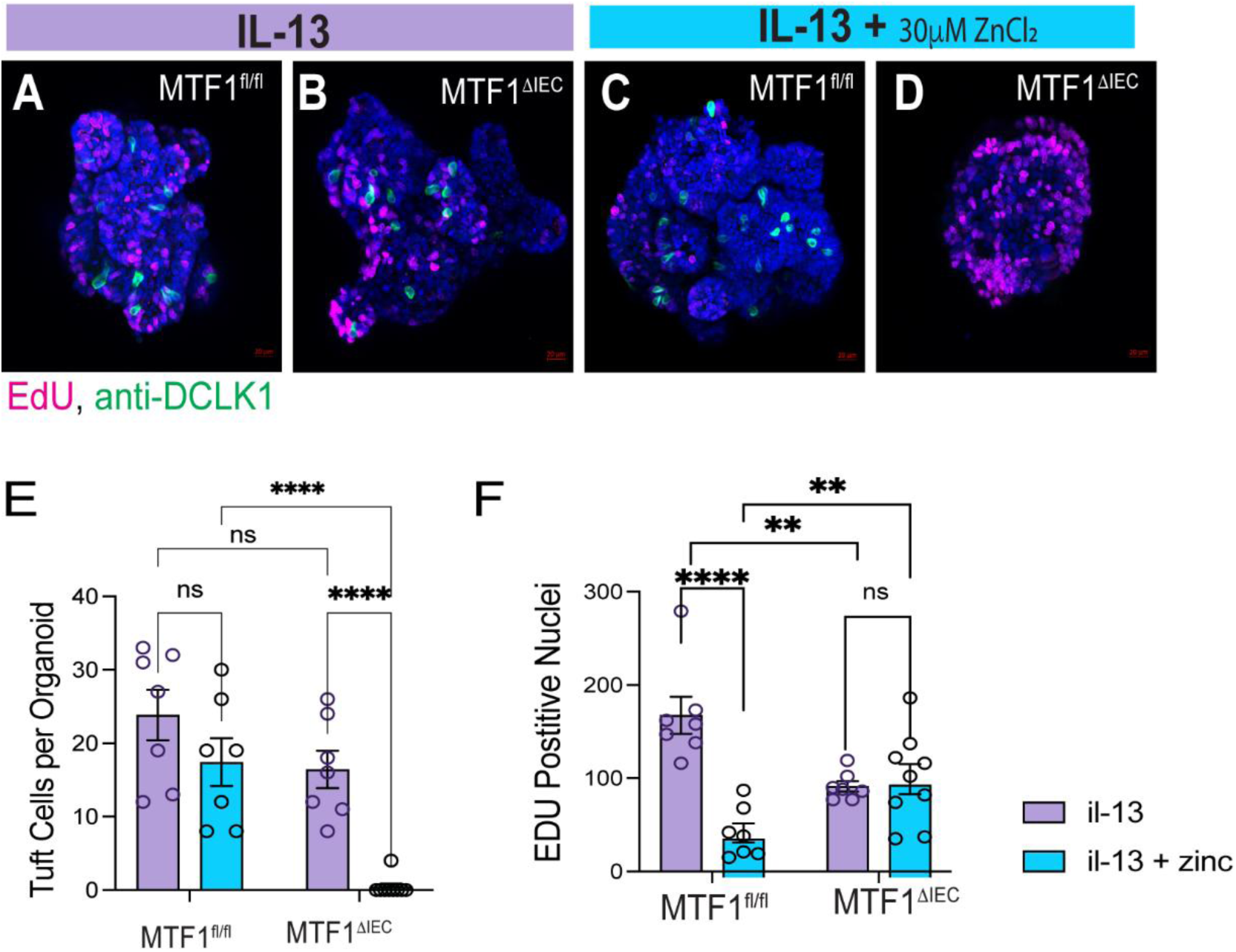
Zn homeostasis via MTF1 is crucial for IL-13 dependent induction of tuft cells ex vivo. Organoids were grown for 7 days with either IL-13 at 10ng/mL alone, or IL-13 and 30 µM ZnCl_2_ present in media. EdU incorporation into SIOs occurred 2 hours before harvest. A-B. Whole Mount immunofluorescent staining of organoids using DCLK1(green) antibody, EdU (Pink) and with DAPI (blue) after IL-13 stimulation. C-D. Same as A-B, with IL-13 and 30µM ZnCl_2_ present in media. E. Number of DCLK1+ cells present in organoids for each condition. F. EdU positive nuclei were counted per organoid, as observed from whole-mount staining for each condition.

## Discussion

Diet and microbiome derived cues are key determinants of epithelial barrier function. In the gut, Zn deficiency has been associated with impaired barrier function and increased intestinal permeability [43, 44]. Several human diseases associated with mutations in Zn homeostasis have intestinal pathogenesis. Acrodermatitis enteropathica is a well-documented inherited condition characterized by Zn deficiency, which is often caused by mutations in the ZIP4 gene (SLC39A4) that encodes Zn transporter and is known to cause Crohn’s like symptoms in carriers [13]. A pleiotropic missense variant in SLC39A8, another Zn transporter gene, has been associated with Crohn’s disease [14]. Additionally, mutations in SLC39A13 have been implicated in a connective tissue disorder that may affect intestinal function [45]. Zn mismanagement resulting from these genetic alterations is thought to lead to impaired mucosal integrity, compromised intestinal barrier function, and altered immune responses in the gut. While the general importance of Zn in intestinal health and barrier function is established, several key molecular and cellular details regarding how Zn sensing in IECs specifically impact the differentiation and function of epithelial barrier are missing. Additionally, how IEC intrinsic Zn homeostasis influence host-microbiome interactions in the gut is not well established.

The current body of work establishes Zn’s importance for epithelial barrier function at both cellular and molecular levels. By utilizing genetic mouse models of constitutive and inducible IEC-specific deletion of MTF-1, a key Zn-sensing transcription factor, our study provides new insights into the critical role of Zn homeostasis in maintaining barrier function *in vivo*. The use of *ex vivo* organoid models further elucidates the cell-intrinsic role of Zn sensing in proliferation and differentiation. We reveal that Zn acts as a trigger for switching from proliferation to differentiation in intestinal epithelial cells, highlighting the intricate relationship between Zn homeostasis and cell fate decisions. By combining *in vivo* and *ex vivo* approaches, this study offers a comprehensive view of how Zn regulates epithelial barrier function through multiple mechanisms, including cellular proliferation, differentiation, and its effect on host-microbe interactions and immune homeostasis via maintenance of key epithelial cell lineages.

Our findings on the simultaneous loss of both Paneth cells and tuft cells in MTF1^ΔIEC^ mice highlight an intriguing connection between these two specialized epithelial cell lineages. Recent studies have shed light on the crosstalk between Paneth cells and tuft cells, suggesting a co-regulatory mechanism that may be influenced by Zn homeostasis. For instance, research has shown that activation of the succinate receptor SUCNR1 in tuft cells can trigger a type-2 immune response, leading to changes in Paneth cell function and promoting dysbiosis and inflammation [36, 37]. Conversely, Paneth cell-derived factors may influence tuft cell differentiation and maintenance [46]. We find that although both tuft and Paneth cells are missing in constitutively deleted MTF1 mice, in the inducible model of MTF 1 deletion, tuft cell defect appears first followed by Paneth cell, suggesting that tuft cell are uniquely sensitive to Zn signals. Additionally we find that the loss of tuft cell lineage ex vivo is replicated but not in the Paneth cell lineage. The absence of both cell types in our *in vivo* mouse model but not *ex vivo* model suggests that Zn-dependent MTF-1 signaling in tuft cells could be a critical upstream regulator of Paneth cell function via tuft cell-induced immune cell circuits. Furthermore, the interplay between these cell types may explain the profound dysbiosis and inflammatory response observed in MTF1^ΔIEC^ mice. The loss of tuft cell-mediated type-2 immune responses, coupled with altered Paneth cell function, likely creates a permissive environment for microbial encroachment and shifts in community composition. This co-regulation hypothesis opens new avenues for understanding how Zn homeostasis orchestrates the delicate balance between different epithelial cell lineages and their collective role in maintaining intestinal homeostasis.

Our findings shed new light on the unique role of Zn homeostasis in tuft cells, underscoring a previously unknown role of dietary micronutrients in regulating host-microbiome symbiosis. Tuft cells, known for their ability to sense a multitude of microbial and metabolic stimuli and generate immune responses that remodel the intestinal epithelium, have recently been discovered to possess an even more remarkable capability [47, 48]. These rare cells can act as reserve stem cells in both humans and mice, giving rise to all other lineages when canonical stem cells are damaged in a IL-13-dependent manner that triggers tuft cell expansion[47]. Our study demonstrates that Zn homeostasis, mediated by MTF-1, is crucial for the differentiation of tuft cells in context of IL-13 stimulation. This finding adds a new layer of complexity to tuft cell biology, suggesting that Zn may play a pivotal role in regulating their stem cell-like properties. The loss of tuft cells in MTF1^ΔIEC^ mice, coupled with their inability to maintain these cells in the presence of Zn in organoid cultures, raises intriguing questions about the role of Zn in modulating tuft cell plasticity and their capacity to act as reserve stem cells. Future studies will focus on elucidating the molecular mechanisms by which Zn regulates tuft cell differentiation, maintenance, and their potential for dedifferentiation into stem cells.

Our findings on the role of Zn in regulating the balance between proliferation and differentiation in intestinal epithelial cells align with and extend previous studies on Zn’s critical function in cell cycle progression. Zn is widely recognized as essential for growth and proliferation, with its deficiency leading to cell cycle arrest [40, 41]. Our *ex vivo* organoid culture results, showing Zn as a trigger for switching from proliferation to differentiation, provide a new perspective on how dietary Zn levels might influence cell fate decisions in the intestinal epithelium. MTF-1, as a Zn-sensing transcription factor, plays a crucial role in maintaining the precise intracellular Zn levels required for fast-dividing cells to progress through all stages of the cell cycle. Previous studies have demonstrated that Zn deficiency or excess can induce quiescence and that Zn resupply or time to bring excess Zn levels down stimulates synchronized cell-cycle reentry [41]. Our observations in MTF1^ΔIEC^ organoids complement these findings by showing that in the absence of Zn sensing, cells maintain a proliferative state at the expense of differentiation. This suggests that MTF-1-mediated Zn homeostasis is essential not only for cell cycle progression but also for the transition from proliferation to differentiation. The Zn-dependent switch we observed in wild-type organoids, where Zn treatment promoted differentiation into specialized cell lineages, can be understood in the context of Zn’s role in cell cycle regulation. It appears that MTF-1 acts as a molecular rheostat, fine-tuning intracellular Zn levels to either promote cell cycle progression or trigger differentiation [41]. In fast-dividing cells, MTF-1 likely maintains Zn levels within a range that supports continuous proliferation. However, when Zn levels rise above a certain threshold, possibly due to environmental cues or cellular signals, MTF-1 may alter its regulatory activity to promote differentiation instead. The inability to properly sense and respond to Zn levels results in a failure to make this transition, leading to continued proliferation even in the presence of Zn in MTF1^ΔIEC^ organoids. These findings highlight the intricate relationship between Zn homeostasis, cell cycle regulation, and differentiation in the intestinal epithelium, underscoring the importance of MTF-1 in coordinating these processes.

In conclusion, our study illuminates the critical role of Zn homeostasis, mediated by MTF-1, in regulating intestinal epithelial function and barrier integrity. By employing both *in vivo* genetic mouse models and *ex vivo* organoid cultures, we have uncovered new insights into how Zn orchestrates the delicate balance between proliferation and differentiation in intestinal epithelial cells. The intricate interplay between Zn homeostasis, epithelial cell fate decisions, and host-microbe interactions revealed in our study opens new avenues for understanding intestinal biology and developing targeted therapies for gastrointestinal disorders. As we continue to unravel the complexities of intestinal epithelial biology, the fundamental role of micronutrients like Zn in shaping tissue function and adaptation becomes increasingly apparent, offering exciting possibilities for advancing both basic science and clinical applications in gastroenterology.

## Resource availability

### Lead contact

Any requests for more information, resources and methodology should be directed to the lead contact, Shipra Vaishnava.

### Materials availability

All strains of mice in this study are available from the lead contact upon request and execution of institutional MTA.

## Acknowledgements

We would like to thank Dr. Samir Beyaz and Onur Eskiocak for their time in helping us begin organoid investigations. S.V. is supported by R01DK113265, P20GM109035 (pilot project) and Brown University Seed grant. R.Y. is supported by T32 HL134625. S.L. is supported by R01 AI183155, R21 AI173821, and the Charles H. Hood Foundation Child Health Research Award. Thermo Apreo VS SEM was purchased using a grant from the Office of the Director at the National Institutes of Health (S10OD023461).

## Author Contributions

Conceptualization, S.V., R.Y.; methodology, S.V., R.Y. G.H. L.Z., S.L.; investigation, S.V., R.Y., G.H., K.B., L.Z.; writing—original draft, S.V. and R.Y.; writing—review and editing, S.V., R.Y., G.H., S.L.; funding acquisition, S.V., S.L.; resources, S.V.; supervision, S.V.

## Methods

### Mice

Mice were bred in barrier facility at the Brown University. Experiments were performed on sex matched mice aged 8-12 weeks, in accordance with protocols approved the Institutional Animal Care and Use Committee. Mice were given access to food and water throughout the day and monitored by staff at Brown University. All mice were littermates or cohoused before the start of experiments, and only separated by sex and treatment during the course of the experiment.

### Generation of MTF1^fl/fl^ mice

To generate MTF1-floxed mice sperm from C57BL/6N-*Mtf1^tm1a(KOMP)Wtsi^* were purchased from the Knockout Mouse Project Repository and the Mouse Biology Program at the University of California, Davis. to fertilize C57BL/6 females. The MTF1-floxed mice have previously been generated and validated for cell specific MTF1 deletion in chondrocytes [49]. Pups carrying *Mtf1^tm1a(KOMP)Wtsi^* allele were crossed to FlpO to remove the FRT-Neo-FRT cassette and generate MTF1-floxed animals. MTF1^fl/fl^ mice were crossed to Villi-cre (Jax# 004586) and Vill-creERT(Jax# 020282) to generate constitutive and inducible MTF1 deletion in the intestinal epithelium.

### Mouse Infections

For *Heligmosomoides polygyrus* infection, 8-12 week old mice were given oral gavage with 200 L3 *H. polygyrus*. Fecal egg burden was tested weekly by counting on McMaster Chamber as previously described and were killed at time indicated for tissues for staining [50]. For MNoV infection, stocks of MNoV strain CR6 were generated from molecular clones as previously described[51]. Adult mice at 8-12 weeks of age were inoculated with 106 PFU of CR6 strain by the oral route in a volume of 25uL. Fecal pellets were harvested on Day 3 and 7. The mice were sacrificed on Day 7 and both stool and tissue samples were flash frozen by dry ice and stored into 2 mL tubes (Sarstedt, Cat#72.694.300) with 1 mm-diameter zirconia/silica beads (Biospec, Cat# 11079110z) at -80°C.

### Induction of Cre-ERT2 by Tamoxifen

MTF1^ERTΔIEC^ and littermate controls were given tamoxifen at 50mg/kg by intraperitoneal injection for 5 days to induce deletion on MTF1 by Cre-ERT2. Mice were observed at 2- and 4-weeks post induction.

### Quantitative Reverse Transcription-PCR for MNoV Infection

RNA was isolated from stool using Quick-RNA Viral 96 kits (Zymo, Cat# R1041) according to the manufacturer’s protocol. 5uL RNA from stool or 1ug of total RNA from tissue was used for cDNA synthesis with ImPromII reverse transcriptase system (Progema, Cat# PAA3803). Quantitative PCR was performed by using AmpliTaq gold DNA polymerase with gold buffer and MgCl_2_ kit (Thermo Fisher Scientific, Cat #4311806). MNoV TaqMan assays were performed, using a standard curve for determination of absolute viral genome copies, as described previously[52]. Standard curves for quantitative PCR assays were used to facilitate absolute quantification of transcript copy numbers of Rps29. Then the absolute values of MNoV per microgram of RNA were normalized to the housekeeping gene Rps29 for tissue samples.

### Lamina Propria and Intestinal Epithelial Lymphocyte Isolation

Mice were euthanized and the terminal half of the small intestine was flushed with ice-cold PBS and cut into small 3-4 cm sections. Sections were washed in cold PBS by vortexing four times, and then the epithelial layer was removed by incubating sections for 30 minutes at 37°C in HBSS (4.17 mM NaHCO_3_, 3%FBS, and 1mM EDTA). To isolate Epithelial Lymphocytes, the supernatant was collected by filtering with a 70uM cell strainer, and then resuspended in 40% Percoll (60% RPMI) to be put on a 40%:80% Percoll gradient. To isolate lamina propria lymphocytes, after HBSS incubation samples are washed again in PBS by vortexing, and then incubated in RPMI (4.17 mM NaHCO_3_, 10%FBS, 0.5X Pen-Strep, 0.5X Na-pyruvate, collagenase, dispase, and DNaseI). This incubation was performed twice for 25 minutes each at 37°C, and each time the supernatant was collected through a 70 uM sterile cell strainer, and was spun at 400gx10 minutes. Cells were then resuspended in 40% Percoll(60% RPMI) and spun on a 40%:80% Percoll gradient. Lymphocytes were collected from the layer between 40% and 80% Percoll, and then cells were stained for intracellular and extracellular markers.

### Flow Cytometry Antibody Staining

Lymphocytes collected from Lamina Propria isolation or Epithelial isolation were resuspended in RPMI (4.17 mM NaHCO_3_, 10%FBS, 0.5X Pen-Strep, 0.5X Na-pyruvate) and incubated for 4 hours at 37°C with Cell Stimulation Cocktail and Protein Transport Inhibitor (eBioscience). After stimulation, cells were stained with fluorescent antibodies that recognized surface antigen for 30 minutes at room temperature in the dark. Cells were washed with dPBS (3% FBS) and then incubated with Fixation/Permeabilization solution (eBiosceinces) overnight at 4°C. Intracellular staining of intracellular cytokines and transcription factors was performed for 30 minutes at room temperature in the dark.

### Metal Analysis

Total Levels of Zn, manganese, iron and copper were measured by Inductively Coupled Plasma OES. To begin, mice were euthanized and the ileum was flushed with ice-cold PBS and cut into small 3-4 cm sections. For whole tissue analysis, one section was weighed and added to a tube for digestion. For analysis of IECs, sections were washed four times in cold PBS by vortexing, and then the epithelial layer was removed by incubating sections for 30 minutes at 37°C in HBSS (4.17 mM NaHCO_3_, 3%FBS, and 1mM EDTA). Supernatant was filtered through 70micron filter, and epithelial cells were pelleted and washed by PBS. Then,100 to 200 mg of tissue was digested in 1 mL of trace metal grade nitric acid for 5 hours at 55°C. Metal-free conical tubes (VWR) were used to ensure no additional metals were added, and nitric acid only mock samples were included for controls. Once intestinal samples were completely dissolved, sterile DI water was used to dilute samples by 1:25, and then they were run on a Thermo Scientific iCAP 7400 DUO ICP-OES.

### Fluorescent In situ Hybridization

Sections of the distal small intestine were preserved in methacarn and embedded into paraffin blocks to make 7uM thick slides for Fluorescent In-Situ Hybridization (FISH). Slides were treated with xylenes and then consecutive ethanol steps until samples were fully hydrated. A 16S RNA probe was used to target bacteria in hybridization buffer of 0.9M NaCl + 20mM Tris-HCl + 0.1% SDS, overnight at 50°C. Slides were washed three times, and then DAPI was used to observe the nuclei of host cells in the SI.

### Scanning Electron Microscopy

Small sections (5-10 mm) of the terminal ileum were fixed in a solution containing 2.5% glutaraldehyde, 0.15 M sodium cacodylate, 2% paraformaldehyde, 2mM calcium chloride, and 0.1M sucrose, with a pH of 7.4. Tissues were incubated for 2 hours at room temperature, and then stored at 4°C. Samples were then washed three times at room temperature in a solution of 0.15 M sodium cacodylate, 2mM calcium chloride, and 0.1M sucrose. Samples were then fixed overnight in 2% Osmium tetroxide, 2.5% glutaraldehyde, 0.15 M sodium cacodylate and 2% paraformaldehyde. After second fixation set, samples were washed in DI water three times, and then dehydrated in an ethanol gradient. Each ethanol step was incubated for 20 minutes at room temperature, and then 100% ethanol was incubated twice. Samples were flash dehydrated using a critical point dryer, and then mounted on coverslips for imaging. Thermo Apreo VS SEM was used for imaging.

### Immunofluorescent Staining of Mouse Tissues

Sections of the distal small intestine were preserved in formalin and embedded into paraffin blocks to make 7uM thick slides for staining. Deparaffinization was done by first treating with xylenes and then progressive ethanol + water washes to hydrate tissues. Slides were immersed in citrate buffer at 95°C for 20 minutes for antigen retrieval. Slides were washed in PBS, then blocked in a 1% BSA + 0.5% Triton X-100 PBS solution for 30 minutes at RT. Slides were washed in fresh PBS, then incubated overnight at 4°C with antibody in 1X PBS + 0.05% Triton X-100. The next day, slides were washed three times in PBS, then secondary antibody was applied for 1 hour at RT. Slides were washed again in PBS, DAPI was stained for 15 minutes at RT, and then coverslips were mounted on, and samples were imaged using an Evident APX 100 or Zeiss Axio Observer Z1 microscope.

### FITC-Dextran Intestinal Permeability Assay

Mice were fasted for 4 hours, then given 6mg/gram bodyweight FITC-dextran 4 kDa by oral gavage, 3 hours before mouse was euthanized. Blood was collected via cardiac puncture and serum was collected from blood by letting it sit undisturbed at room temperature for at least 30 minutes. Samples were centrifuged for 10 minutes at 2000 rpm, and serum was collected in sterile tubes. FD4 fluorescence was quantified in duplicates by plate reader (SpectraMax M3, Molecular Devices) using 485/535 nm excitation/emission maxima. Standard curve was made using control serum and FD4.

### Establishment and Care of Organoid Culture

Mice were euthanized at 8-10 weeks and 3 cm sections of the ileum were washed with ice cold PBS until the supernatant was clear. This protocol was developed based on previously published protocols [42]. Briefly, sections were incubated 0.5mM EDTA in PBS, rocking, on ice for 45 minutes. After incubation, samples were washed with fresh PBS, and then consecutive fractions in sterile dPBS were filtered through a 70-micron cell strainer. The cleanest fraction was selected, and then isolated crypts were counted and embedded in Matrigel (Cat#356231, Corning Growth Factor Reduced). Crypts were resuspended in 75% Matrigel and 25% media at approximately 5-10 crypts per uL of media. Matrigel domes were allowed to solidify for 10 minutes at 37°C and then growth media was added to each well. Growth media was comprised of Advanced DMEM (Cat# 12634010, Gibco), CHIR-99021 5 μM (Cat# 4423, Tocris), N-acetyl-L-cysteine 1 μM (Sigma-Aldrich), Y-27632 10 μM (Cat# 1254, Tocris), N2 1X (Cat# 17502048, Gibco), B27 1X (Cat# 17504044, Gibco), 1% Penicillin-Streptomycin (Cat# P4333, Sigma-Aldrich), 1% GlutaMAX (Cat# 35050061, Gibco). To stimulate growth and budding of organoids, media was supplemented with recombinant murine EGF 40 ng/ml (Cat # 315-09, PeproTech), recombinant murine Noggin 50 ng/ml (Cat # 250-38, PeproTech), and R-spondin 62.5 ng/ml (Cat# 3474-RS, R&D Systems). Growth media was made fresh weekly, and growth factors were added immediately before administering to each well. Cultures were maintained in a fully humidified incubator at 5% CO2 and 37°C. Domes were split twice a week for maintenance, approximately every 3-4 days, and growth media was changed every other day. In all experiments organoids grew to maturity and were harvested on day 7.

### Harvesting and Immunofluorescent Staining of Organoids

Organoids were grown for a period of 7 days with any treatments. For harvesting, media was replaced in each well with 5% EDTA in cold sterile PBS, and plate was placed on cold block. Plate was rocked for 30 minutes, until Matrigel began to dissolve. Each well was harvested separately, and organoids were pelleted by centrifugation for 1 minute at 200 g. Organoids were washed twice by removing supernatant and replacing with fresh cold PBS. Organoids were fixed in 4% PFA for 30 minutes at room temperature, or 45 minutes on ice. PFA was removed after organoids were pelleted, and samples were washed twice with cold PBS. To begin staining, organoids were permeabilized by incubation in a 0.1M Glycin + 0.3% Triton X-100 solution for 10 minutes. Organoids were washed twice in PBS, and primary antibody was added overnight in a solution of 1X PBS + 1% BSA + 0.5% Triton X-100. The next day, organoids washed twice in PBS and then secondary antibody was applied in a solution of 1X PBS + 0.05% Triton X-100. The final day, organoids were washed twice in PBS, then stained with DAPI for 15 minutes at room temperature, then mounted on slides using Prolong Gold Antifade mountant (Invitrogen). Organoid slides were imaged on Zeiss LSM 800 Confocal microscope.

### Edu Labelling of Organoids

After 7 days of growth, organoids were incubated with 10µM EdU in media solution for 2 hours. Organoids were then harvested as previously described. After fixation and washing, the Click-iT™ Plus EdU Cell Proliferation Kit (Invitrogen) was used to label EdU. Briefly, organoids were permeabilized as previously described, washed twice with PBS. A staining solution was made composed of 1X Click-iT Buffer, copper protectant, component B, and 1X Click-iT buffer additive which was administered to the organoids and incubated at room temperature for 30 minutes in the dark. Organoids were then washed twice in PBS, and Hoechst 33342 stain was used to label nuclei. After two washes with PBS, organoids were then mounted on slides in Prolong Gold Antifade mountant (Invitrogen) and imaged on Zeiss LSM 800 Confocal microscope.

### Quantitative Real-Time-PCR Tissues

Whole tissue RNA was extracted from terminal ileum using PureLink™ RNA Mini Kit (Invitrogen) and cDNA was synthesized using M-MLV Reverse Transcriptase (ThermoFisher). SYBR green master mix (Thermo Fisher Scientific) was used to perform qPCR. Gene expression was normalized to the housekeeping gene, GAPDH. Changes in gene expression were displayed relative to controls and calculated using the DD CT method for analysis.

### Quantitative real-time-PCR Organoids

After 7 days of growth organoids were harvested from Matrigel domes by incubation with 5mM EDTA in 1X PBS for 30 minutes on ice. Organoids were centrifuged for 1 minutes at 200g, then supernatant was removed. Organoids were washed twice in 1X PBS, then RNA was extracted using PureLink™ RNA Mini Kit (Invitrogen). Organoid cDNA was synthesized (M-MLV Reverse Trascriptase, ThermoFisher). SYBR green (ThermoFisher) was used for qPCR, and the DD CT method was used for analysis. Gene expression changes were normalized to GAPDH, and fold changes were relative to controls.

#### 16S Sequencing and Microbiome analysis

##### gDNA Extraction and quantitative Real-Time-PCR of Microbiome

Genomic DNA extraction was performed on fecal, luminal, and mucosal samples using the Quick-DNA Fecal/Soil Microbe Microprep Kit (Zymo Research). To collect luminal samples, a 1mL sterile PBS flush was collected from a portion of the ileum, spun by centrifugation at 5,000rpm for 10 minutes and then supernatant was removed. To collect mucosal scraping, after luminal flush intestines were filleted open and scraped into 1mL of sterile PBS. Mucus sample was spun at 5,000rpm for 10 minutes and then supernatant was removed. For qPCR analysis of microbiome, 16S primers were used to observe total bacterial burden of SI luminal contents, and probes specific to T. muris were used to observe burden in the feces.

##### 16S rRNA gene sequencing and analysis

The V4/V5 region of the bacterial 16S rRNA gene was amplified using the Phusion High-Fidelity DNA polymerase (Thermo Scientific) with 518F/926R primer pair (518F: 5’- CCAGCAGCYGCGGTAAN -3’, 926R: 5’- CCGTCAATTCNTTTRAGT -3’), which were conjugated with overhang adapter sequences for Illumina MiSeq. The amplification program consisted of 95 °C for 3 min, followed by 35 cycles of 95 °C for 30 sec, 58 °C for 30 sec, and 72 °C for 30 sec, and 72 °C for 10 min. DNA libraries were constructed and sequenced on the Illumina MiSeq platform (paired-end 300 bp sequencing) at Rhode Island Genomics and Sequencing Center.

The raw sequencing reads were processed using the QIIME2 pipeline [53]. The raw sequencing reads were trimmed and denoised followed by amplicon sequence variants (ASVs) table construction using DADA2 [54]. The pre-trained Naïve Bayes classifier on the SILVA 132 database [55] was used for taxonomic assignment. Singletons and all features annotated as mitochondria or chloroplast were removed from the table. The relative abundance of bacterial taxa was expressed as a percentage of total 16S rRNA gene sequences.

##### RNA-Seq analysis

Extracted RNA samples were submitted to the GENEWIZ or Novogene for library construction and sequencing. The RNA library was prepared using NEBNext Ultra RNA Library Prep Kit for Illumina (New England Biolabs) and sequenced on Illumina HiSeq or Illumina NovaSeq X Plus platform (Paired-end 150 bp).

Adapter and low-quality sequences were trimmed from the raw sequence read using Trimmomatic version 0.36 [56]. Trimmed sequences were aligned to the mm10 mouse genome using STAR v2.7.3a [57]. Differentially expressed genes (DEGs) between groups were identified if Padj < 0.05 in the R package DESeq2 v1.38.1 [58]. Gene set enrichment analysis (GSEA) was performed and visualized using the R package clusterProfiler v4.7.1 [59]. For GSEA, a pre-ranked gene list based on log2FoldChange was used as an input, and the lists of cell marker genes were obtained from PanglaoDB [25]. Heatmap was drawn using the R package ComplexHeatmap v2.15.2 [60].

### Data code and availability

Raw sequence reads from 16S sequencing have been deposited in the NCBI Sequence Read Archive (SRA) under accession number PRJNA1198599. Bulk RNA-seq data has been deposited in the NCBI GEO database under accession numbers GSE284340 and GSE284341.

### Statistical Analysis

Data were analyzed with R software and Prism 10 software, and data was expressed as ±SEM. Significance was determined by unpaired two-tailed student’s t-test between two groups, if data followed normal distribution. If data did not follow normal distribution, Mann-Whitney test was used to compare two groups. For multiple group comparison, one-way ANOVA was used. The n value details can be found in figure legends. In the graphs, * indicates a significant p value of <0.05, ** indicates a very significant p value of <0.01, and *** indicates a highly significant p value of <0.05.

---

**Figure S1:**
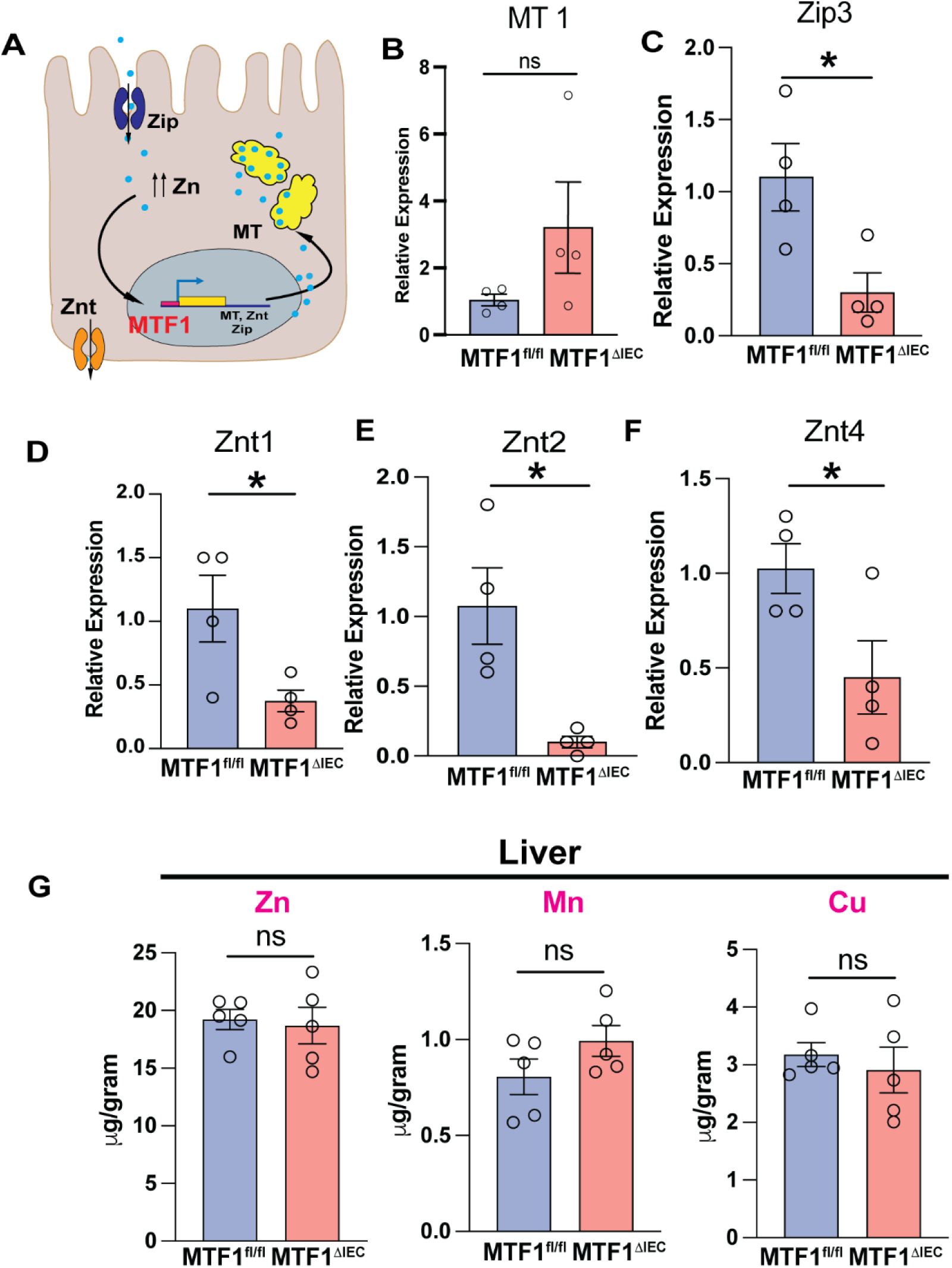
Loss of Zn-sensitive MTF1 results in altered expression of Zn-machinery. A. Schematic diagram of the role MTF1 plays in IEC. Zn transport proteins (ZIP) bring Zn into IEC, which increases cytoplasmic Zn levels. MTF1 senses excess Zn and increases transcription of various Zn-related machinery such as metallothioneins (MT), ZIPs and Zn export proteins (ZnT). Metallothioneins bind free cytoplasmic Zn, serving as an intracellular Zn reservoir. B-F. Metallothionein-1, Zn Importer Protein 3, and various Zn Transport protein expression by qPCR (n=4). G. ICP-OES total quantification of metals (bound and free) of whole tissue liver, displaying levels of zinc, manganese and copper in MTF^ΔIEC^ compared to littermate controls(n=4).

**Figure S2:**
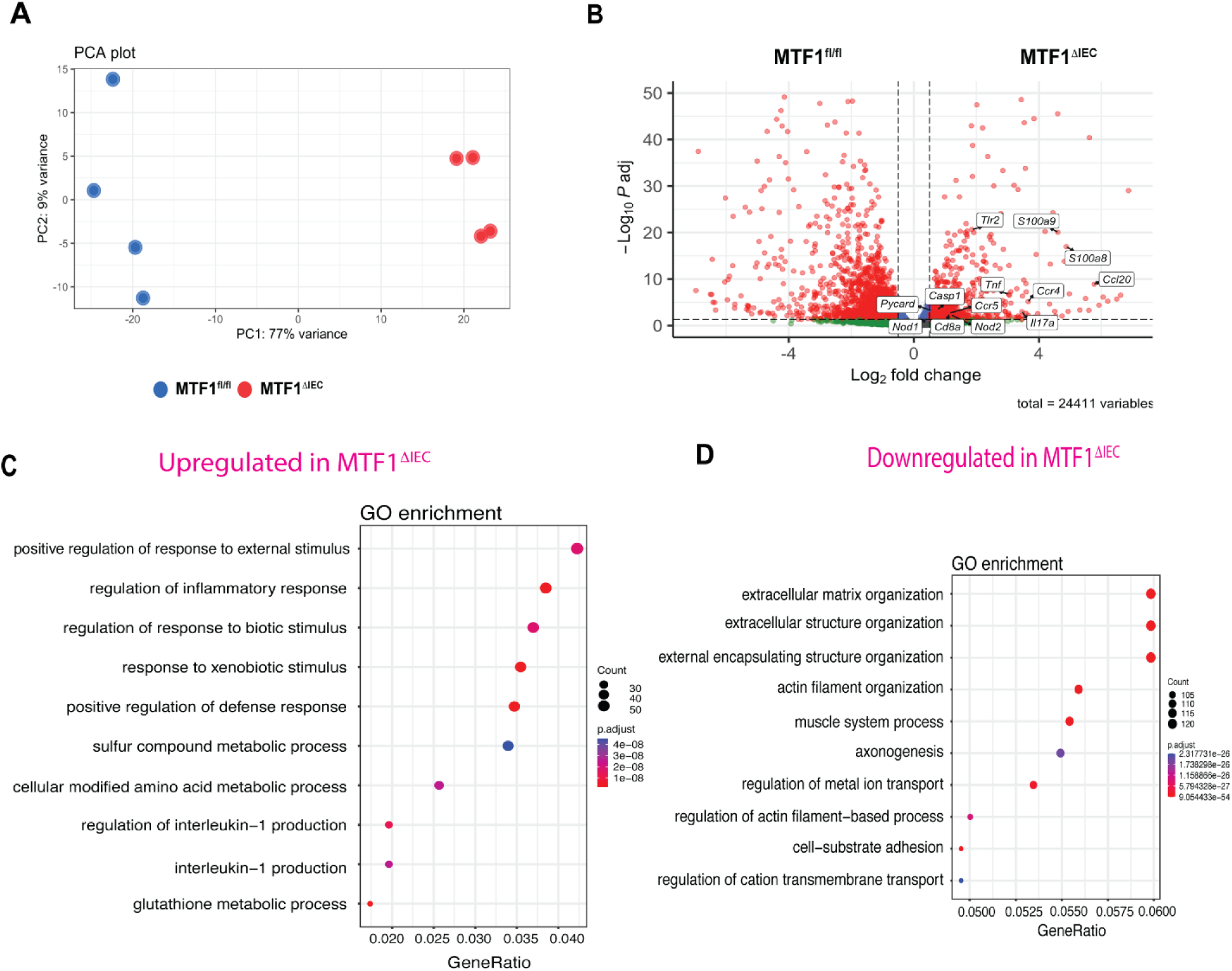
Ileum of MTF1^ΔIEC^ showed distinct transcriptional profiles by bulk RNA-sequencing. A. Principal component analysis plot shows a clear cluster separation in homeostasis ileum tissue from MTF1^fl/fl^ and MTF^ΔIEC^. (n=4) B. Volcano plot of differentially expressed genes between MTF1^fl/fl^ and MTF1^ΔIEC^ mice. C. Gene Ontology (GO) plot showing genes upregulated in MTF1^ΔIEC^ mice compared to control. D. GO plot showing genes downregulated in MTF1^ΔIEC^ mice compared to control.

**Figure S3:**
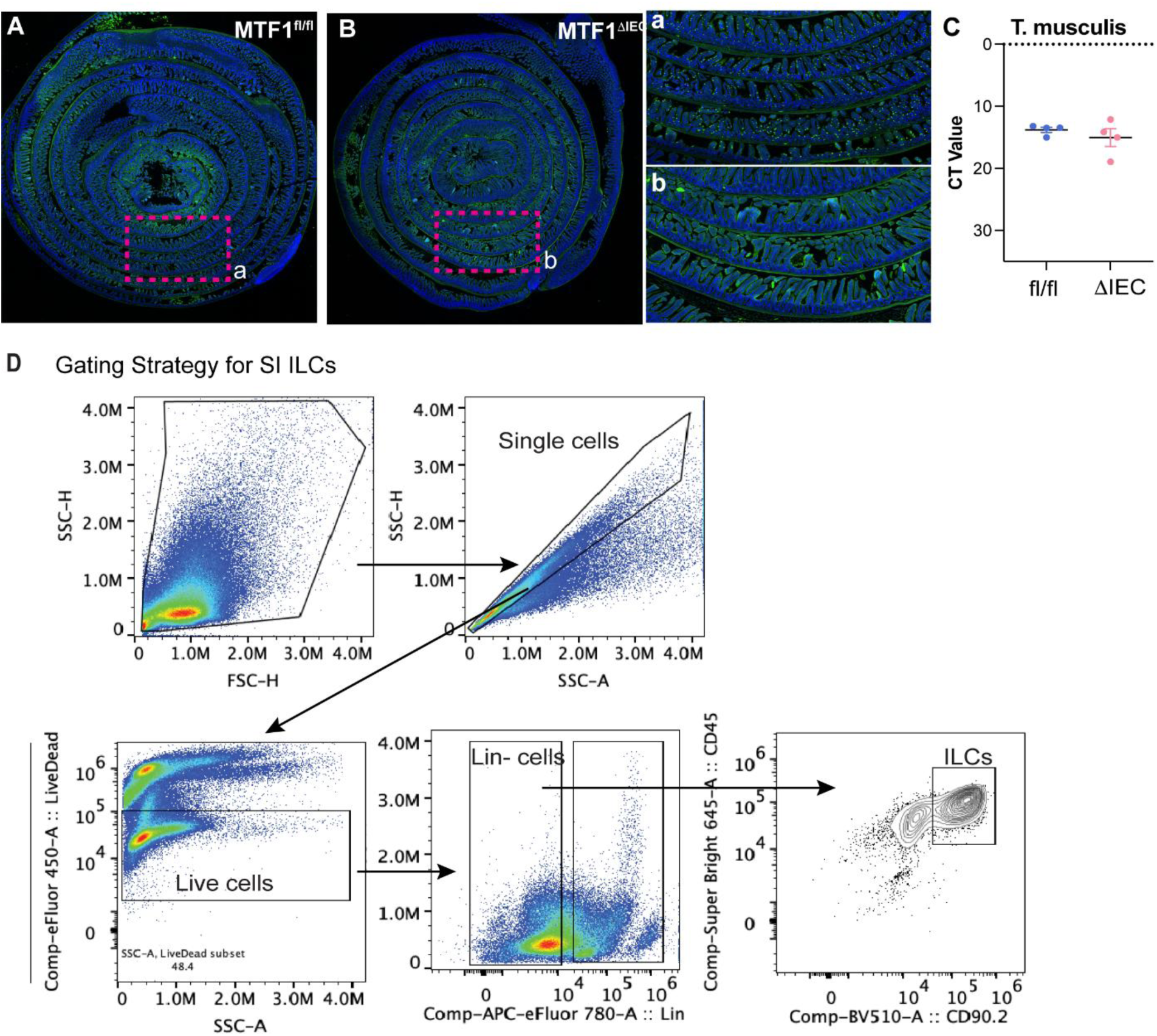
Loss of tuft cells in MTF1^ΔIEC^ is consistent across SI and is independent to tuft cell-stimulating microbes. A-B. Tuft cells (green) by immunofluorescent staining of DCLKM-1 (DCLK-1) and DAPI (blue) of seen by swissroll of jejunum. C. *Tritrichomonas musculus* load seen by qPCR of feces. (n=4) D. Gating strategy of flow cytometry analysis for observing ILC populations from epithelial fractions of SI.

**Figure S4:**
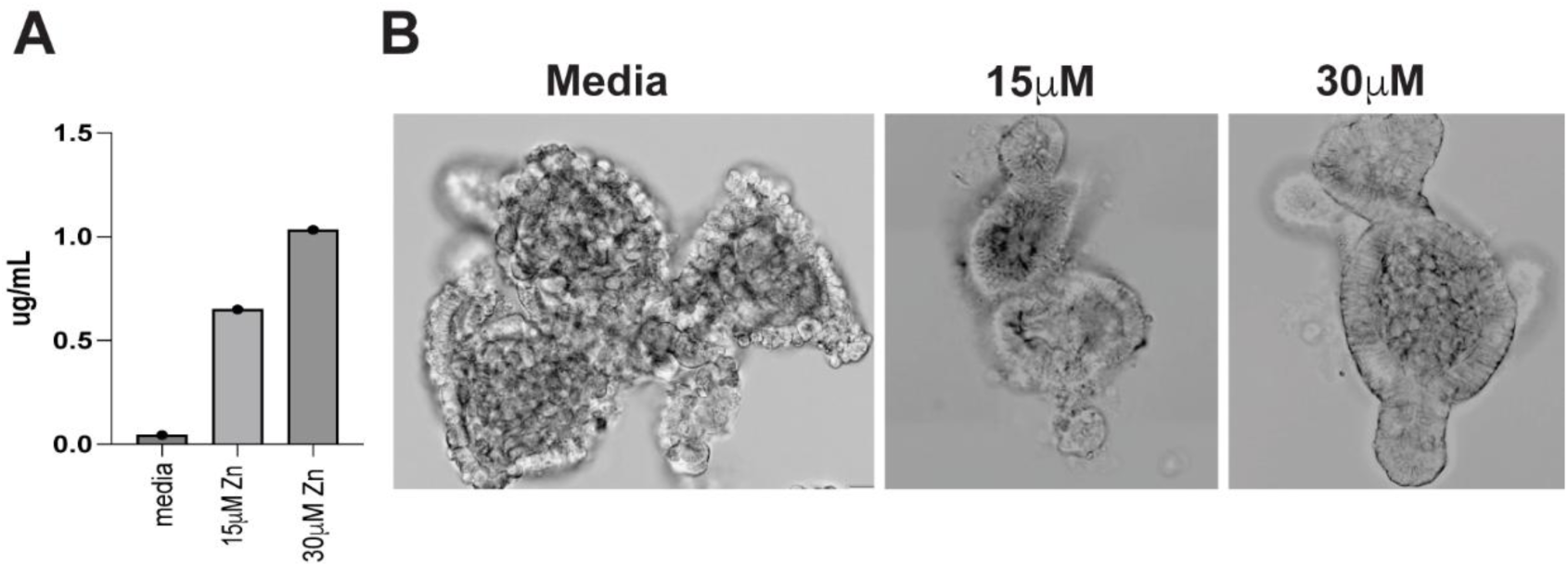
Zinc quantification in organoid media and its impact on organoid morphology. A. ICP-OES results of Zn levels present in media, showing control, 15 µM and 30 µM treatments of ZnCl_2_. B. Brightfield whole-mount MTF1^fl/fl^ organoids showing development of budding and typical characteristics in control, 15µ M and 30 µM treatments of ZnCl_2_.

**Figure S5:**
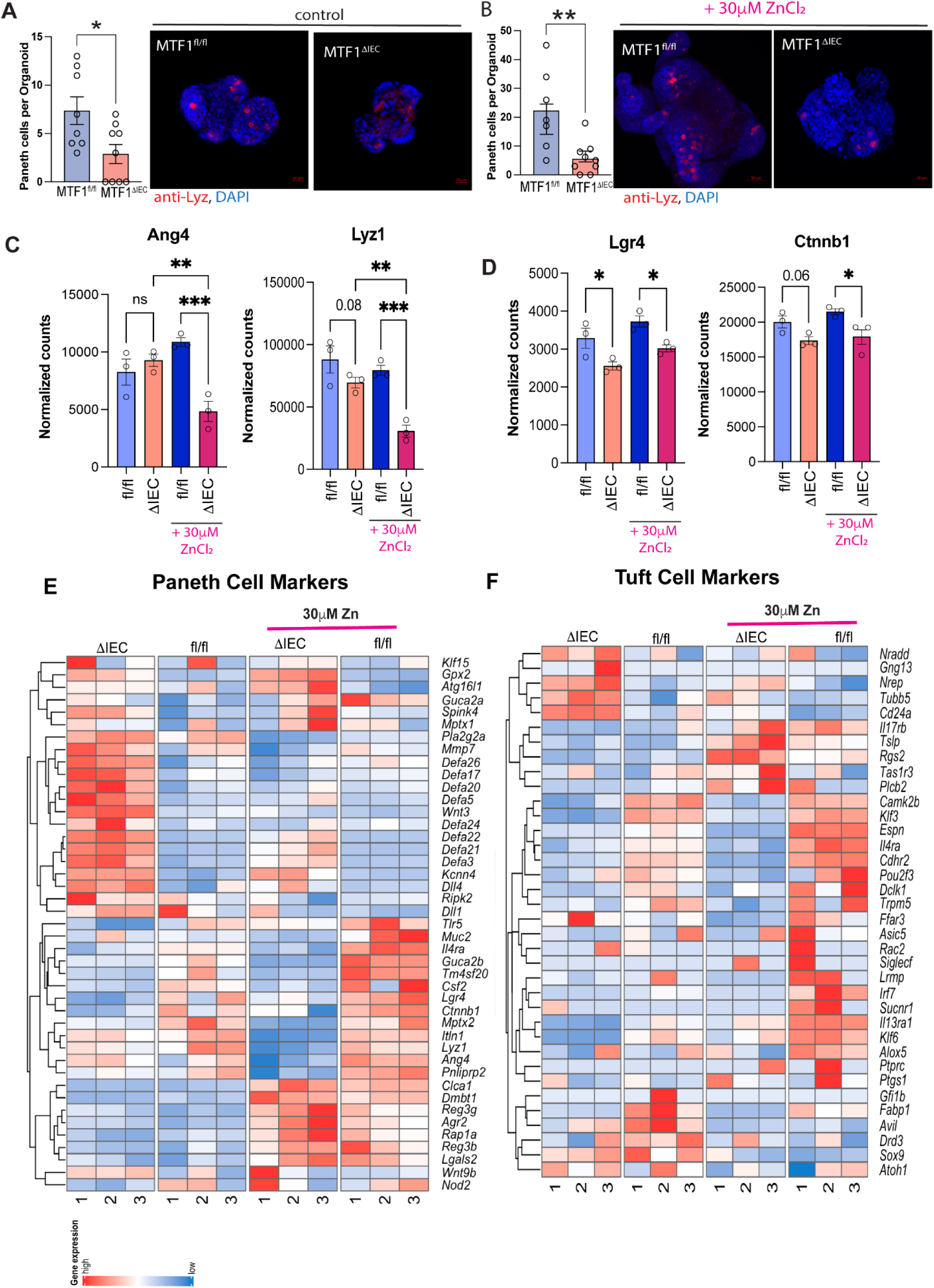
Paneth cell in MTF1^fl/fl^ and MTF1^ΔIEC^ SIOs without and with Zn treatment. A. Whole-Mount Immunofluorescent staining of organoids using lysozyme (red) and DAPI (blue) during homeostasis (Zn-free) conditions. Number of lysozyme producing cells were counted per organoid. B. Same as A, with 30µM ZnCl_2_ present in media. C. Paneth cell antimicrobial markers present in SIOs with or without 30µM ZnCl_2_ treatment (n=3). D. Stem cell markers present in SIOs with or without 30µM ZnCl_2_ treatment (n=3). Significance was determined by DESeq2 analysis. E. Heatmap of gene expressions related to tuft cells from RNA-sequencing of MTF1^ΔIEC^ and MTF1^fl/fl^ SIOs with or without 30µM treatment of ZnCl_2_ in media (n=3 wells per condition). F. Same as E, but for tuft cell markers.

